# New exon ignites accelerated evolution of placental gene *Nrk* in the ancestral lineage of eutherians

**DOI:** 10.1101/2021.02.09.430412

**Authors:** Guopeng Liu, Chunxiao Zhang, Yuting Wang, Guangyi Dai, Shu-Qun Liu, Wenshuai Wang, Yi-Hsuan Pan, Jianping Ding, Haipeng Li

## Abstract

Accelerated evolution is often driven by the interaction between environmental factors and genes. However, it remains unclear whether accelerated evolution can be ignited. Here, we focused on adaptive events during the emergence of chorioallantoic placenta. We scanned the chromosome X and identified eight accelerated regions in the ancestral lineage of eutherian mammals. Five of these regions (*P* = 5.61 × 10^−11^ ~ 9.03 × 10^−8^) are located in the five exons of Nik-related kinase (*Nrk*), which is essential in placenta development and fetoplacental induction of labor. Moreover, a eutherian-specific exogenous exon lack of splice variant was found to be conserved. Structure modelling of NRK suggests that the accelerated exons and the eutherian-specific exon could change the enzymatic activity of eutherian NRK. Since the eutherian-specific exon was surrounded by accelerated exons, it indicates that the accelerated evolution of *Nrk* may be ignited by the emergence of the new exon in the ancestral lineage of eutherian mammals. The new exon might shift the function of *Nrk* and provide a new fitness landscape for eutherian species to explore. Although multiple exons were accelerated in both of the *Nrk* catalytic and regulatory domains, positive selection can only be revealed on the regulatory domain if the branch specific nonsynonymous and synonymous rate test was performed by PAML. Thus, it may be important to detect accelerated evolution when studying positive selection on coding regions. Overall, this work suggests that the fundamental process of placental development and fetoplacental induction of labor has been targeted by positive Darwinian selection. Identifying positively selected placental genes provides insights into how eutherian mammals gain benefits from the invasive chorioallantoic placenta to form one of the most successful groups among terrestrial vertebrates.

## Introduction

Eutherian mammals (more commonly referred to as placental mammals) form one of the most successful groups among terrestrial vertebrates. The success is mainly due to the fact that they have evolved a most invasive chorioallantoic placenta. The placenta provides a direct contact of the fetal tissue with the maternal blood [1–4], and hence an intimate association between the fetus and the maternal system.

The placenta is the first organ to form during mammalian embryogenesis. It acts as the interface between the fetal and the maternal environments, and thus allows an exchange of gases, nutrients and waste products between the mother and the fetus [5]. Fetal membranes are considered as prerequisites for amniotes, the most successful land vertebrates, to reproduce independently from aquatic environments. It has been proposed that the formation of fetal membranes triggers the independent evolution of highly complex placentas in marsupial and eutherian mammals [3]. Although many genes have been reported to be associated with the development of placenta [5–7], the questions of how the eutherian has evolved the chorioallantoic placenta [8–10] and which genes underwent positive Darwinian selection during the emergence of chorioallantoic placenta [11], remain unanswered. Moreover, it may provide insights into whether accelerated evolution can be ignited.

A previous study was carried out to screen the autosomal regions that have undergone an accelerated evolution in the ancestral lineage of eutherians [12]; however, the identified accelerated regions in the autosomes are unlikely to be related to the development and function of placenta since their nearest genes do not highly express in the placenta. The X chromosome (chrX) was not investigated because the evolution rate of chrX is lower than that of the autosomes [13]. In this study, we screened the chrX to detect the regions with accelerated evolution in the ancestral lineage of eutherians. Out of the eight significantly accelerated regions on the chrX, five regions, including the most accelerated one (Figure 1, and Table S1), are located in the exons of Nik-related kinase (*Nrk*) [14,15]. NRK, which belongs to the germinal center kinase-IV (GCK-IV) subfamily within ste20-type family of mammalian protein kinases [16], plays a key role in placental development and fetoplacental induction of labor [7].

**Figure 1.**
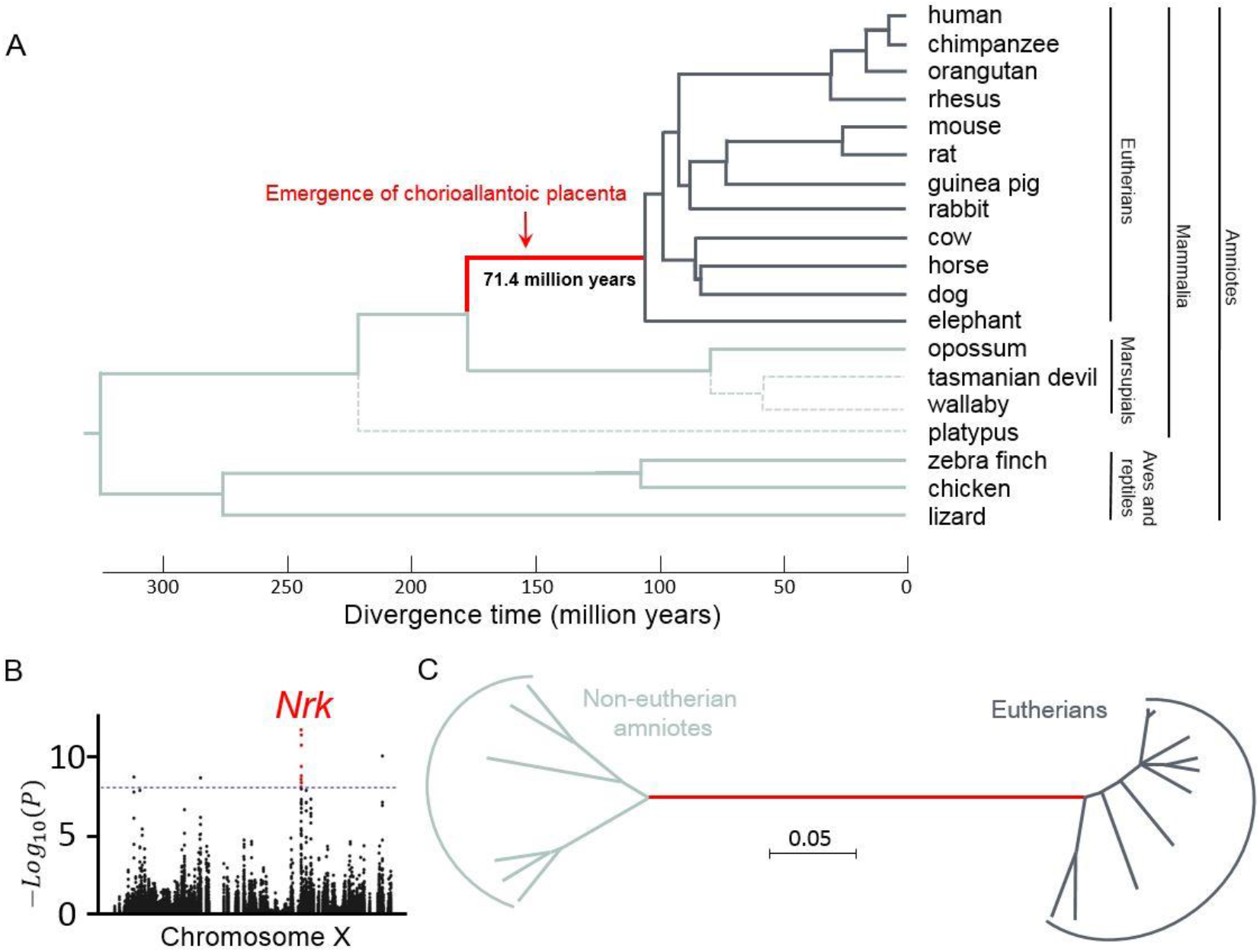
Genome-wide screening for placental-accelerated sequences on chromosome X. (A) Phylogenetic tree of 19 amniotes for detecting placental-accelerated sequences. The red branch indicates that chorioallantoic placenta has emerged in the ancestral lineage of eutherian mammals. Sixteen amniotes linked by solid lines were used in the Kung-Fu Panda (KFP) test for accelerated evolution. Three other amniotes (tasmanian devil, wallaby, and platypus) linked by dashed lines were included to validate the results. (B) Manhattan plot of KFP screening results. Results of genome-wide scan for sequences with accelerated evolution are shown in Manhattan plot of significance against human chromosomal locations. Each dot represents one window. The location of the windows with the highest signal is indicated in red. The dash line denotes the threshold of the test after Bonferroni correction. (C) Unrooted Neighbor-Joining tree of the conjunction of 5 accelerated regions located in *Nrk* gene. Except for lizard, sequences of all species showed in (A) were used. The evolutionary distances were computed using the Tajima-Nei method [48].

Interestingly, by estimating the branch specific nonsynonymous to synonymous rate ratio, and examining different regions of *Nrk* gene, PAML (Phylogenetic Analysis by Maximum Likelihood) [17] failed to reveal the positive selection in the most of examined cases (4/5). It may explain why the accelerated evolution of placental-related *Nrk* gene has never been found before. The protein structure modelling reveals that the accelerated and eutherian-specific exons encoding the catalytic domain of NRK are likely to participate in the regulation of its enzymatic activity. This work suggests that the new exon ignites the accelerated evolution of placental gene *Nrk*, and the fundamental process of placental development and fetoplacental induction of labor has been targeted by positive Darwinian selection.

## Results

A phylogenetic tree of 16 amniotes (12 placental mammals, 1 marsupial, 2 aves, and 1 reptile) was constructed, and the length of each branch was determined as in the previous study [12] (Figure 1A). Then, using the Kung-Fu Panda (KFP) software [12], we screened the chrX to detect the regions with accelerated evolution in the ancestral lineage of eutherians. A total of 99,106 windows were scanned, and 13 of them were identified (*P* < 1.01 × 10^−7^) (Figure 1B) after Bonferroni correction [18] for multiple tests (the desired overall Type I error rate, *i.e.* the family-wise error rate, is 0.01). The overlapped windows were clustered into eight different regions, which were defined as chrX Placental-Accelerated Sequences (chrX-PASs, Table S1). Notably, five of the eight chrX-PASs, including the most accelerated one (*P* = 5.61 × 10^−11^), are located in the coding regions of *Nrk*. NRK is a member of the GCK subfamily which regulates various developmental processes and MAPK (Mammalian mitogen-activated protein kinase) core signaling pathways [19,20]. Specifically, NRK has been shown to play an important role in the regulation of placental development and fetoplacental labor induction [7].

To validate the accelerated regions, their syntenic alignments were visually examined. It was observed that sequence substitutions in these accelerated regions might have happened between the clades of placental mammals and marsupials-aves. Moreover, the neighbor-joining tree of *Nrk* shows that *Nrk* is not (highly) conserved within the placental and non-placental clades (Figure 1C). A very long red branch confirms that *Nrk* has experienced the accelerated evolution in the ancestral lineage of eutherians.

It is of interest to compare the accelerated regions on the autosomes with those on the chrX. Only six out of 28 accelerated autosomal windows were located in coding regions [12], while seven out of eight accelerated chrX regions in coding regions. Despite this large difference (21.4% *vs* 87.5%), it may not indicate the preference of selection between non-coding and coding regions. Considering that 11 out of 28 autosomal accelerated windows were located in the non-coding social enhancer PAS1 [12], and five out of eight accelerated chrX regions were located in *Nrk*, the differed contribution of coding regions might be explained by the clusters of accelerated regions.

Since the coding regions of eutherian *Nrk* has experienced the accelerated evolution, we then used the PAML package [17] to detect the lifted ratio between nonsynonymous and synonymous substitution rates (ω) of the most accelerated exon 5 (Table S1). However, PAML failed to identify the positive selection on *Nrk* exon 5. The branch-specific model results showed that the ω values for the ancestral lineage of eutherians and other branches were 0.3227 and 0.036, respectively. Moreover, the ratio of nonsynonymous to synonymous substitution rates was calculated for the exon 5 of human and Tasmanian devil using DnaSP [21]. The ω value is 0.183, which also failed to reveal positive selection on *Nrk*. Similar conclusions were made when considering the other accelerated exons, the catalytic domain and conjunction of the catalytic domain and the regulatory domain (Table S3). It was found that the gene was positively selected only when its regulatory domain was considered (ω = 1.122). Therefore, it seems that the Kung-Fu Panda is more sensitive for the signature of positive selection on coding regions than the ω-related methods, and thus can provide a refined analysis to narrow down selected regions.

Next, the alternative hypothesis was explored for the accelerated evolution of *Nrk*. Loss-of-function could cause accelerated evolution [22,23], but it is impossible in our case as *Nrk* still functions. It is also unlikely that the accelerated evolution of *Nrk* could be explained by GC-biased gene conversion [24]. Such an explanation needs to assume that recombination hotspots occurred on the five exons only in the ancestral lineage of placental mammals; this is, however, an un-supported scenario. Thus, the accelerated evolution of *Nrk* is very likely driven by positive Darwinian selection.

As a kinase, NRK is composed of a catalytic domain (CD) and a regulatory domain. The CD of NRK plays a critical role in the AKT (protein kinase B) phosphorylation [25]. The *Nrk* gene in human is composed of 10 exons (exon 2-11, Figure 2B). Despite the large difference in the amino acid sequences of the CDs between eutherians and non-eutherian amniotes, many of the amino acid substitutions come from the exon 5 (Figure 2A). Since directly identifying the potentially beneficial functions of those substitutions is challenging, an alternative way is taken to predict whether the ancestral allele is (slightly) deleterious, instead of predicting whether the derived allele is (slightly) beneficial. This was accomplished by using the PROVEAN (Protein Variation Effect Analyzer) test [26]. The results show that, when compared with the human alleles, the non-eutherian alleles in the exon 5 may encode deleterious amino acids: 85N, 118L and 119S, due to their low PROVEAN scores of −3.98, −8.05 and −5.73, respectively. These results are in accordance with the assumption that *Nrk* has undergone accelerated evolution driven by beneficial functional changes during the emergence of eutherians.

**Figure 2.**
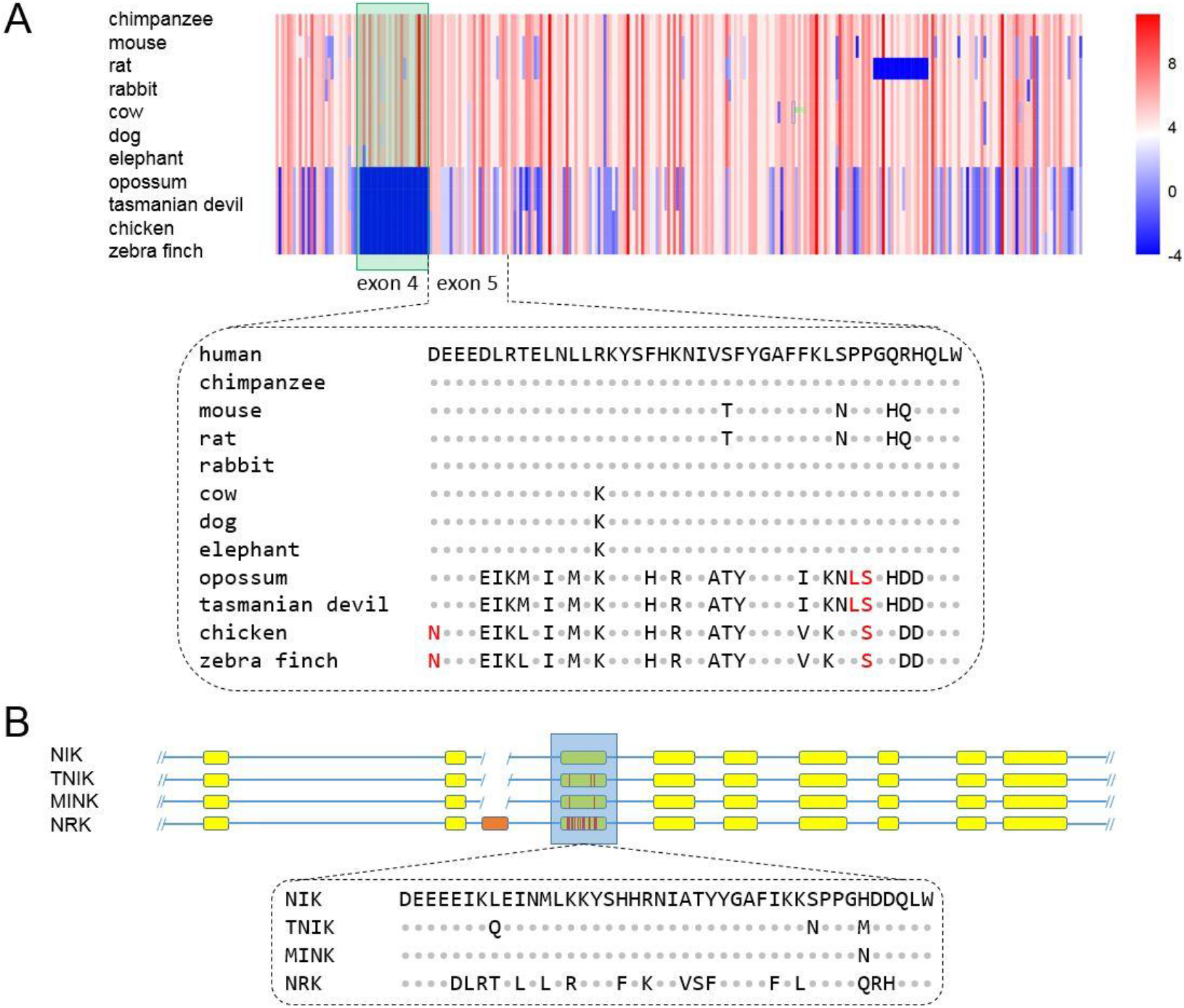
Accelerated evolution of eutherian NRK. (A) The amino acid sequence identity when comparing human NRK to eutherians (mouse, rabbit, dog and elephant) and non-eutherian amniotes (opossum, tasmanian devil, chicken and zebra finch) orthologs. The score was calculated from the amino acid substitution matrix [52]. Human NRK exon 4 (green box) has no ortholog in non-eutherian amniotes, so the score value was assigned as −4. Dotted block shows the sequences of human NRK exon 5 and its orthologs. To human, the amino acids substitutions colored red are predicted by PROVEAN to be deleterious. (B) Catalytic regions of human GCK-IV subfamily. Human NRK exon 4 is indicated in orange. Sequences of human NRK exon 5 and its paralogs are shown in dotted block.

Interestingly, when compared to marsupials and aves, eutherians have a eutherian-specific exon 4 (Figure S1). It did not emerge until the common ancestor of eutherians emerged, and it has been well conserved since its origin (Figure S1). This is different from the cases of new genes, which in their early stage usually underwent rapid changes in sequence and continued an evolution of function [27]. Furthermore, we performed a BLAST search and found no evidence of exonization of intronic sequence [28] or duplication of other exons. It indicates that the eutherian-specific exon did not generate from the endogenous DNA material. We also searched the NCBI viral sequence database but did not found orthologs of the exon 4. Unlike most newly born exons with splice variants, all transcripts of eutherian *Nrk* contain the eutherian-specific exon 4. Thus the exon is not spliced during mRNA maturing, which suggests that it may play an important role in the development. This is even more striking when considering the overall similar structures of the CDs of GCK-IV subfamily members (Figure 2B). As shown, the CD of eutherian NRK contains the eutherian-specific exon 4, while the structures of the CDs of the other GCK-IV subfamily members remain the same.

Further analysis demonstrated that the eutherian-specific exon is surrounded by accelerated regions (Figure S1). The exon 4 is neighbored by the most accelerated region (exon 5). Since the exon 3 encodes only 19 amino acids, it is unsuitable for detecting accelerated evolution. Therefore, the exons 2 and 3 were joined together to be subjected to accelerated evolution detection (*P* = 1.69 × 10^−7^). Therefore, it could be suggested that the emergence of the eutherian-specific exon is likely to ignite the accelerated evolution of *Nrk* in the ancestral lineage of eutherian mammals.

To further study the function of the accelerated exon 5 and the conserved eutherian-specific exon 4, 3D structure models of the NRK-CD were constructed using the homology modelling technique based on the crystal structures of the CDs of Traf2- and Nck-interacting kinase (TNIK) and mitogen-activated protein kinase kinase kinase kinase 4 (MAP4K4) (Figure 3A; Table S2). In comparison with the inactive forms of TNIK-CD and MAP4K4-CD, the α1-helix in the active forms is shifted to the ‘in’ conformation (Figure 3B). Superposition of the NRK-CD homology models of three species reveals a similar situation. Moreover, the loop 4 (L4; residues 61-84, encoded by the eutherian-specific exon 4 in human NRK, *h*NRK) is located outside of the α1-helix, which is the most accelerated region encoded by the exon 5, indicating a potential function (Figures 3C,D, and 4).

**Figure 3.**
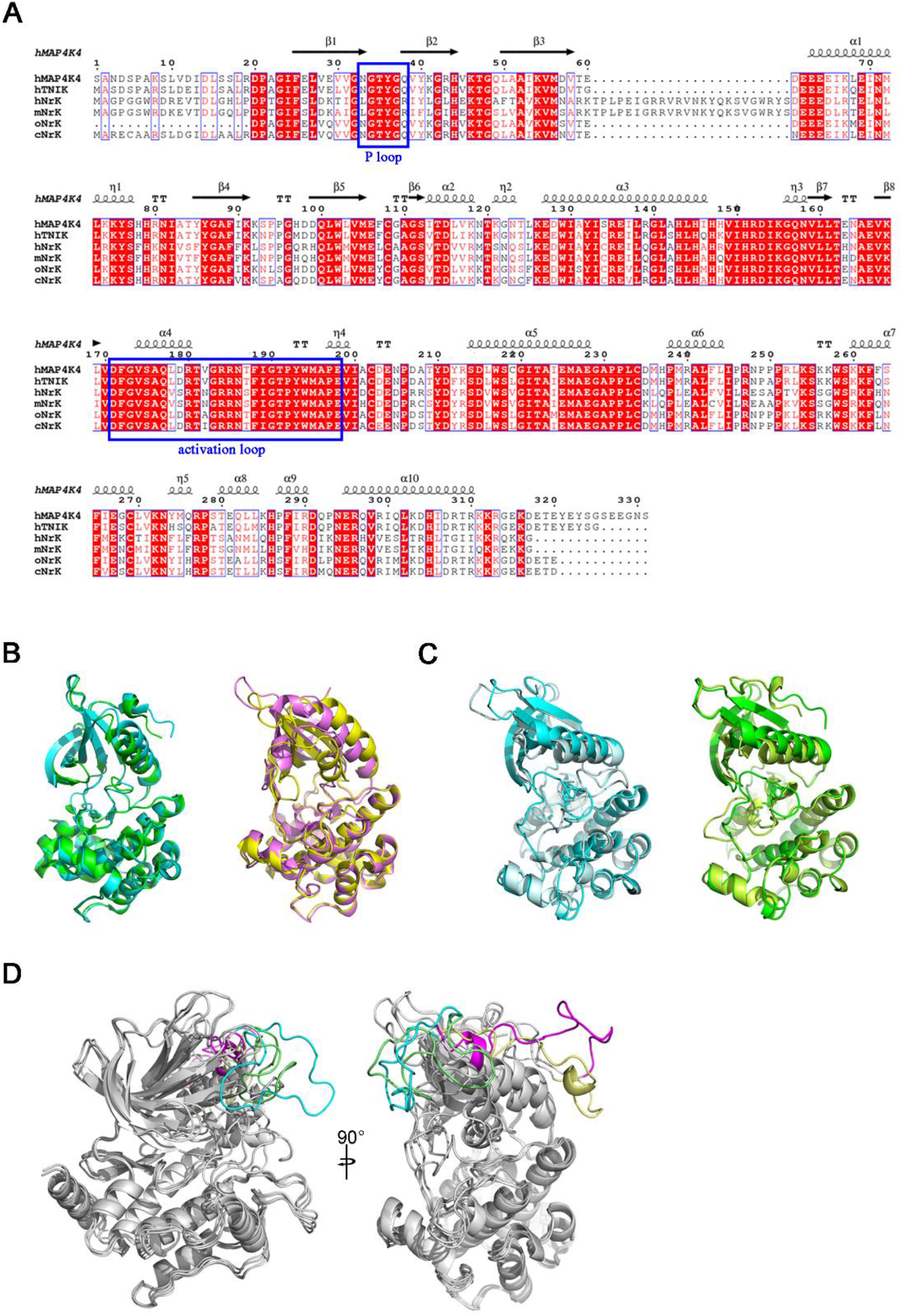
Structural and sequence analyses of TNIK, MAP4K4 and NRK. (A) Sequence alignment of MAP4K4-CD, TNIK-CD and NRK-CD. The lower letters before the gene name represent different species (h, m, o and c represent human, mouse, opossum and chicken, respectively). (B) Left panel: superposition of active TNIK (green, PDB code: 5ax9) and inactive TNIK (cyan, PDB code: 5cwz). Right panel: superposition of active MAP4K4 (magenta, PDB code: 4u40) and inactive MAP4K4 (yellow, PDB code: 4u3y). (C) Superposition of TNIK-templated *o*NRK (left) and *c*NRK (right) in different conformations. Active forms are shown in palecyan and limon. Inactive forms are shown in cyan and green. (D) Different orientations of the L4 region in the homology models. The L4 regions of active and inactive TNIK-templated (PDB codes: 5ax9 and 5cwz) models are highlighted with lime and cyan, respectively. The L4 regions of active and inactive MAP4K4-templated (PDB codes: 4u40 and 4u3y) models are highlighted with magenta and yellow, respectively.

**Figure 4.**
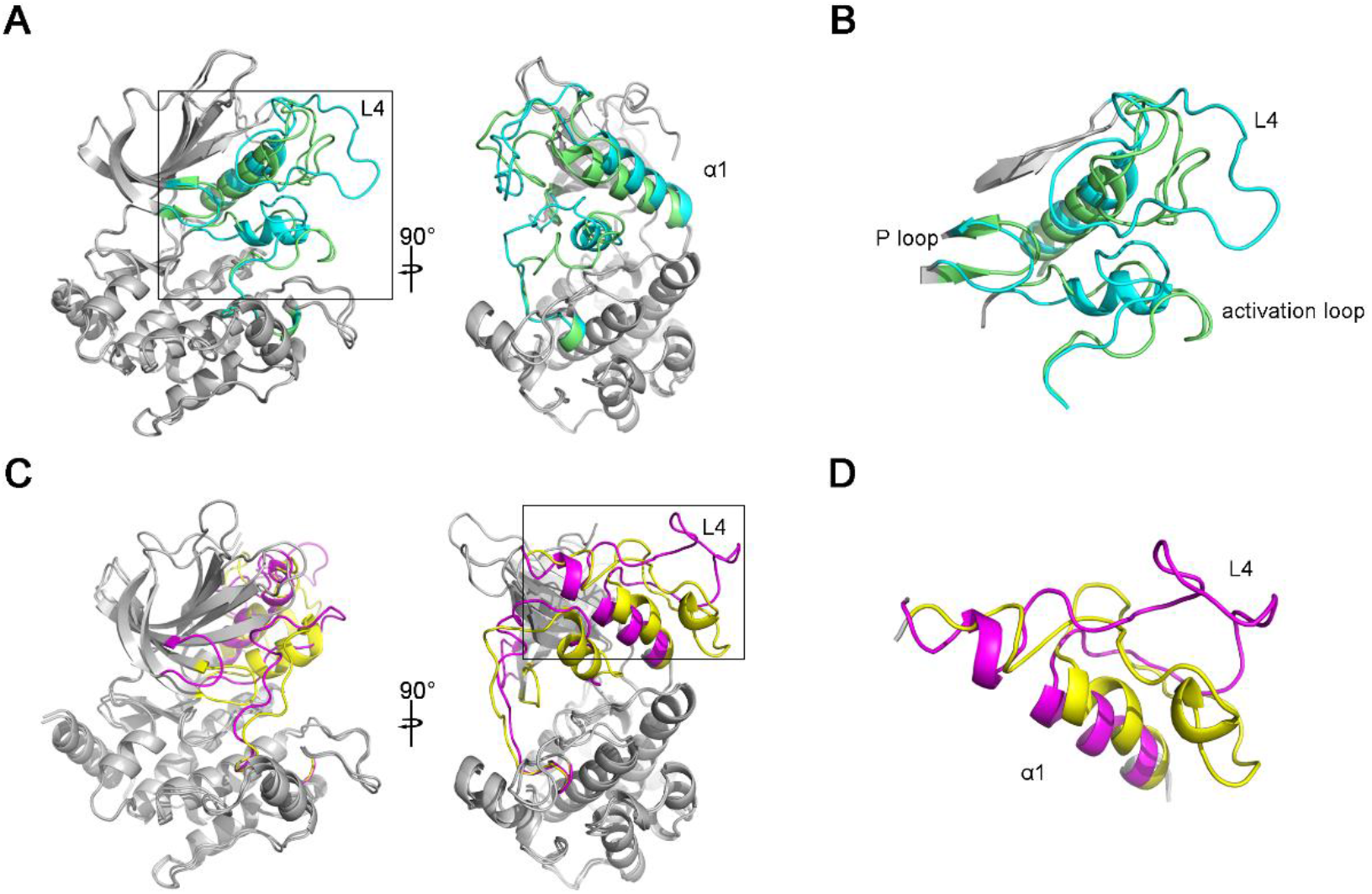
Homology models of *h* NRK-CD and the potential function of the L4-α1 region. (A) Comparison of the TNIK-templated homology models of active and inactive *h*NRK-CDs. PDB codes for the active and inactive TNIK template are 5ax9 and 5cwz, respectively. The active-site regions in the two models are highlighted with lime and cyan, respectively. (B) Detailed view of the active-site regions composted of the L4, α1, P-loop, and activation loop. Color coding is the same as in (A). (C) Comparison of the MAP4K4-templated homology models of active and inactive *h*NRK-CDs. PDB codes for the active and inactive MAP4K4 are 4u40 and 4u3y, respectively. The active-site regions in the two models are highlighted with magenta and yellow, respectively. (C) Detailed view of the L4-α1 region in (C).

As shown in Figure 3D, the L4 demonstrates large differences in the conformations and orientations among the four structures of (in)active TNIK-templated and (in)active MAP4K4-templated models, reflecting the high flexibility of this loop. Of note is that although the L4 is not close to the P-loop and activation loop in the NRK primary amino acid sequences (Figure 3A), in the TNIK-templated *h*NRK models, it is positioned in close proximity to the P-loop and activation loop. In addition, the P-loop, also known as the glycine-rich loop, which is encoded by the accelerated exons 2 and 3 (Figure S1), remains highly conserved and plays a role in the interaction with ATP in the catalytic process [29,30]. The activation loop, which has the regulatory phosphorylation site or interacts with activity modulators, is defined as the segment between two conserved motifs (DFG…APE). In the inactive state, the activation loop collapses in the active center and hinders the substrate binding [31,32]. Thus, it seems likely that the L4 could participate in regulating the catalysis-related conformational changes and/or stabilizing the ‘in’ form of the α1-helix (Figure 4A, B) via its interactions with the other two functionally important loops. In the MAP4K4-templated models, the L4 is an insertion at the α1-helix N-terminus, and therefore its high flexibility may further disrupt the integrity of the α1-helix (Figure 4C, D). In either of the two cases, it may indicate a new mechanism that the eutherian-specific L4 (exon 4) of NRK could act together with the α1-helix (the accelerated exon 5) to modulate the enzymatic activity. Thus, both the accelerated exon and the conserved eutherian-specific exon of *Nrk* might have contributed collectively to the emergence of fully functional chorioallantoic placenta in the eutherian lineage.

## Discussion

*Nrk* gene was first cloned from mouse embryos, and *Nrk* expression could be detected only in the embryonic stage rather than the adult stage [14,33], suggesting that it may be involved in the developmental process of mouse embryos [7]. *Nrk* knockout mice fetus showed expanded placenta and irregular spongiotrophoblast layer, a fetus-derived region of placenta, and pregnant mice exhibited a severe defect in delivery when all fetuses were homozygous mutant ones [7]. In mouse, the placenta is composed of three layers: maternal decidua, junctional zone and labyrinth [34]. The spongiotrophoblast is part of the junctional zone and its loss and expansion can cause the embryonic lethality [35] and the overgrowth of fetus and abnormal delivery [7,36]. Thus, *Nrk* plays a key role in placental development and fetoplacental induction of labor [7].

In this study, counting exon 2 and 3 on, we found that there are seven exons in *Nrk* that have experienced accelerated evolution and identified one eutherian-specific exon in the ancestral lineage of eutherians. Labor induction occurs in the late gestation and is strictly controlled to make the offspring delivered safely. Marsupials give birth at a very early stage of development and complete their growth in the pouch for a long period of lactation [37,38]. For eutherians, it has been found that the labor-inducing signals transmitted from the fetus or the placenta to the mother [7,39]. Moreover, uterine transfer of *Nrk*-null fetuses to wild-type females revealed that pregnant mice exhibit a severe defect in delivery [7]. Therefore, the positive Darwinian selection on *Nrk* may be associated with labor-inducing fetoplacental signal.

To date, most identified novel exons in eutherian mammals generated endogenously by whole gene or single exon duplications and exonization mechanisms [28], which means there are paralogs of these new exons [40–42]. To reduce the disruption of the proteins functions caused by the sudden insertion of new exons, most of these novel exons are alternatively spliced at a much higher frequency than older ones [40,43,44]. In this study, the eutherian-specific exon 4 was found, and there are no alternative spiced variants for this exon. Moreover, no paralogs of the exon were identified in eutherians and non-eutherians, indicating that the exon is generated exogenously. Therefore, the new exon may penetrate the fitness barrier, shift the function of *Nrk*, and bring ancestral eutherians into a new fitness parameter space. Then the adaptation finally resulted in the emergence of chorioallantoic placenta.

There are 28 kinases in mammalian genomes sharing similarity to the budding yeast kinase Ste20p [16]. Among these kinases, Nck-interacting kinase (NIK), Mishappen/NIK-related kinase (Mink), Traf and Nck-interacting kinase (TNIK), and NIK-related kinase (NRK) belong to the germinal center-like kinases IV (GCK-IV) subfamily [20]. More recently, the role of NRK in guaranteeing the normal placental development was attributed to its ability to properly regulate the AKT phosphorylation level [25]. Specifically, the AKT phosphorylation at Ser473 was upregulated in *Nrk*-null trophoblasts, thus resulting in enhanced proliferation of spongiotrophoblast [25]. Our constructed 3D structure models show that the two structural regions, the α1-helix and L4 regions encoded by the most accelerated exon 5 and the eutherian-specific exon 4, respectively, are likely to play a role in regulating the NRK enzymatic activity due to their close locations to each other and the high flexibility of L4. Therefore, the positive selection might have played an essential role in the emergence of chorioallantoic placenta.

Lactation is an important part of mammalian and mammary gland is essential for mammals to take diversified lactation strategies to accommodate reproductive success and to adapt to the environment [37]. In mice, deficiency of *Nrk* during pregnancy triggers mammary gland hyperplasia and adenoma [29]. Considering the evolution history of mammary gland [37] and *Nrk* gene, it is possible that the origin and divergence of lactation are associated to the generation and mutations of *Nrk*.

In summary, this study reveals an important evolutionary process for how the chorioallantoic placenta and the fetoplacental induction of labor have emerged during evolution. It is likely that positive Darwinian selection has favored the conserved eutherian-specific exon of *Nrk*. Since the structural L4 region encoded by the eutherian-specific exon can affect the active-site conformation of NRK, accelerated evolution of the other exons is beneficial to fine-tuning the function of NRK. As the outcome, the *Nrk* gene has gained its new function to modulate placental development and fetoplacental induction of labor.

## Materials and Methods

### Multi-genome alignments

Multi-genome alignments of following 16 representative species of amniotes were created by MULTIZ [45] (http://www.bx.psu.edu/miller_lab/): *Homo sapiens* (human, hg19), *Pan troglodytes* (chimpanzee, panTro3), *Pongo abelii* (orangutan, ponAbe2), *Macaca mulatta* (rhesus, rheMac2), *Mus musculus* (mouse, mm9), *Rattus norvegicus* (rat, rn4), *Cavia porcellus* (guinea pig, cavPor3), *Oryctolagus cuniculus* (rabbit, oryCun2), *Bos taurus* (cow, bosTau6), *Equus caballus* (horse, equCab2), *Canis lupus familiaris* (dog, canFam2), *Loxodonta africana* (elephant, loxAfr3), *Monodelphis domestica* (opossum, monDom5), *Gallus gallus* (chicken, galGal3), *Taeniopygia guttata* (zebra finch, taeGut1), and *Anolis carolinensis* (lizard, anoCar2). Genome sequences and paired genome alignments with human genome as the reference were downloaded from the UCSC genome browser website (http://hgdownload.soe.ucsc.edu/downloads.html).

### Detection of accelerated sequences

The multi-genome alignments were scanned by Kung-Fu Panda (KFP) software [12]. The window size and sliding step were set as 100bp and 20bp, respectively. The Newick tree with branch length from the TimeTree website (www.timetree.org) [46] is ((((((((human:6.4,chimpanzee:6.4):9.3,orangutan:15.7):13.9,rhesus:29.6):61.4,(((mouse:25.2,rat:25.2):46.9,guineaPig:72.1):14.3,rabbit:86.4):4.6):6.4,(cow:84.6,(horse:82.5,do g:82.5):2.1):12.8):7.3,elephant:104.7):71.4,opossum:176.1):148.4,((chicken:106.4,zebraFinch:106.4):168.5,lizard:274.9):49.6):0, where the branch length is in unit of one million years.

### Neighbor-Joining tree

To visualize the accelerated evolution of NRK, the unrooted Neighbor-Joining tree [47] of the conjunction of the 5 exons (*i.e.* exon 5, exon 22 and exons 26-28) which contains accelerated regions was constructed by the eGPS software [30]. The evolutionary distances were computed using the Tajima-Nei method [48] and are in the units of the number of base substitutions per site.

### PAML test

Branch model of PAMLX software [49] was used. The phylogenetic is ((((((human: 6.4, chimpanzee: 6.4): 84.6, ((mouse: 25.2, rat: 25.2): 61.2, rabbit: 86.4): 4.6): 6.4, (cow: 84.6, dog: 84.6): 12.8): 7.3, elephant: 104.7)#1: 71.4, (opossum: 76, tasmanian: 76): 100.1): 148.4, (chicken: 106.4, zebraFinch: 106.4): 218.1):0.0).

### PROVEAN (Protein Variation Effect Analyzer) test

PVOVEAN test was performed online (http://provean.jcvi.org/seq_submit.php). Human NRK sequence was inputted as protein sequence. The difference amino acids between non-eutherian amniotes and human were used as amino acid variations.

### Homology modeling

The modeling server SWISS-MODEL (https://www.swissmodel.expasy.org/) [50,51] was used to build the 3D structure models of *h*NRK-CD, *o*NRK-CD and *c*NRK-CD. The crystal structures of the catalytic domains of human TNIK-CD (PDB codes: 5cwz and 5ax9) [47] and MAP4K4-CD (PDB codes: 4u3y and 4u40) which share the highest sequence identity with NRK were selected as the templates for the homology modeling. Structural analyses and comparisons were performed with PyMOL (http://www.pymol.org).

## Data availability

Genome sequences underlying this article were downloaded from the UCSC genome browser. All relevant data are within the manuscript and its Supporting Information files.

## Authors’ Contribution

**Guopeng Liu:** Evolutionary analysis, writing. **Chunxiao Zhang:** Homology modeling and structure analysis. **Yuting Wang:** Evolutionary analysis. **Guangyi Dai:** Evolutionary analysis. **Shu-Qun Liu:** Analysis, revision. **Wenshuai Wang:** Pre-analysis. **Yi-Hsuan Pan:** Evolutionary analysis, writing, revision. **Jianping Ding:** Supervision, project administration, revision. **Haipeng Li:** Supervision, project administration, revision. All authors read and approved the final manuscript.

## Acknowledgement

This work was supported by grants from the National Natural Science Foundation of China (nos. 31100273, 91731304), and National Key Research and Development Project (No. 2020YFC0847000).

